# Deconstructing the association between abiotic factors and species assemblages in the global ocean microbiome

**DOI:** 10.1101/2020.03.12.989426

**Authors:** Xiaoyu Shan, Otto X. Cordero

**Affiliations:** Department of Civil and Environmental Engineering, MIT, Cambridge, USA

## Abstract

Recent work on microbial communities from various environments has shown that coexisting microorganisms with similar metabolic functions can be combined into high level “functional groups”, which explain a larger proportion of variance in abiotic parameters than any individual taxon. However, the general rules by which taxa should be aggregated into functional groups remain elusive. Here, we show that two conditions are required for species-assemblages to explain a higher percentage of variance in abiotic factors than single taxa. 1) consistent taxa-environment correlations, and 2) weak or negative correlations (i.e. complementarity) between taxa. Applying this recipe to the ocean microbiome, we found that the best grouping of taxa is one that partitions it into only two groups, a core and a flexible assemblage. The core assemblage is enriched in *Cyanobacteria* and oligotrophic heterotrophs, and was strongly correlated to the first principal component of the taxa-sample matrix. The flexible assemblage instead was enriched in metabolically versatile copiotrophs, abundant at higher depths. This simple core / flexible bipartition explained the most variance in abiotic parameters and outperformed annotation-based functional groups as well as individual taxa. It therefore represents the simplest and best grouping of taxa that can be extracted from current ocean microbiome surveys.

## Introduction

Microbial communities often involve thousands of different taxa despite their sharing of many metabolic pathways^1–3^. This observation has led to the notion that the high taxa diversity of a community can be compressed into a much smaller set of functional groups, each encompassing taxa with similar functional profiles^4,5^. However, identifying these functional units in a systematic manner remains an unsolved and challenging problem. Previous research has relied on gene annotations to show that the fractional abundance of metabolic functions is much less variable across biological replicates and across time than that of taxa (typically defined as 16S rRNA ribotypes)^4,6–8^. This suggests that functional groups and taxa are controlled by different ecological pressures^1^. Functional groups, on the one hand, may be strongly coupled with external environmental factors, such as temperature, pH and nutrient concentration^1,2,4,5^. Taxa, on the other hand, may be primarily controlled by community ecology processes such as predation^9^ and interference competition^10^. This implies that the identification of functional groups is an essential step in any effort to map microbial community structure to function and to disentangle the impact of metabolic reactions from the variation induced by community-level processes. However, the challenge remains to learn how to discover the best way to aggregate taxa abundances into relevant functional groups. The availability of large-scale sequencing datasets associated with environmental metadata makes ocean microbiomes a good candidate to address this challenge^11–14^.

A recent study based on the Tara Oceans expedition data showed that, compared to taxa (operational taxonomic units, or OTUs), aggregations of taxa with common metabolic annotations explained a much higher percentage of the variance in abiotic factors such as temperature, nitrate concentration and depth^5^. Therefore, functional groups may not only be less variable than individual taxa, but also better predictors of (and better predicted by) the environment, as expected from the notion that they are primarily controlled by abiotic conditions. Perhaps the best example of a tight association between an environmental parameter and an aggregation of taxa is the correlation between temperature and the first principal component (PC1) of the taxa-sample matrix in the Tara Oceans dataset^11^. PC1 for surface samples and temperature have been found to display a linear regression R^2^ of 0.76^11^, reinforcing the notion that abiotic factors control the abundance and dynamics of large slices of the community that encompass multiple taxa. This result also suggests that functional groups may help us reveal fundamental abiotic controls of community structure. However, it is unclear whether the variation in PC1 captures the variation of a bona fide functional group, given the lack of ecological interpretability of principal components.

In this paper we aim to address two related questions: 1) What properties should an assemblage of taxa fulfill so that the sum of their relative abundance is better correlated to an environmental parameter and less variable than the individual taxa? 2) Is there enough signal in the data that would make it possible to identify functional groups in an unsupervised manner, that is, without resorting to external functional annotation which carry their own biases and limitations? Answering these questions is critical to any effort aimed at identifying functional groups in an annotation-independent manner. In this study we will show what are the general properties required by a group of taxa to display all the desired statistical properties of a functional group when aggregated. In the process, we will reveal the biological interpretation of the PC1 in the Tara Oceans data. Moreover, we will show that, based on the Tara Oceans dataset, the ocean microbiome can be partitioned into two components, a ‘core’ assemblage of microbes, composed mainly of surface associated phototrophs and oligotrophs, and a ‘flexible’ assemblage, enriched in organisms abundant in deeper waters and likely associated to particles. Finally, we will show that taxonomic groups (aggregations of taxa based on their taxonomy at the level of genus, family, etc.) are unlikely to fulfill the statistical properties commonly associated with functional groups (stability and linkage to abiotic factors).

## Results

### Partitioning the ocean’s microbiome into core and flexible assemblages of taxa

In order to explore the fundamental structure of the subtropical ocean microbiome we started by asking whether there was evidence for the existence of a ‘core’ assemblage of taxa, widely represented across the different sampling stations of the Tara expedition. For each taxon (OTUs extracted from the metagenomic data^5,11^), we quantified its occupancy frequency – i.e., the percentage of samples in which the taxa was present. We found that the distribution of occupancy frequency values was u-shaped, with a large peak near zero and a smaller peak near 100% (Fig. 1a). U-shaped distributions are a universal feature of pangenomes (where the analysis is based on genes instead of taxa) and even technological systems such as software projects made of multiple interconnected components^15^. The u-shaped occupancy frequency distribution of taxa gives us an objective way to partition the microbiome set into a ‘core’ and a ‘flexible’ assemblage, by separating those taxa with occupancy frequency higher or lower than a critical point of the frequency histogram. We calculated the first derivative of the smoothed frequency function and found a critical point around 70%, where a sudden bump in the first derivative indicated the emergence of the second peak. (Fig. 1a & Fig. S1).

**Figure 1.**
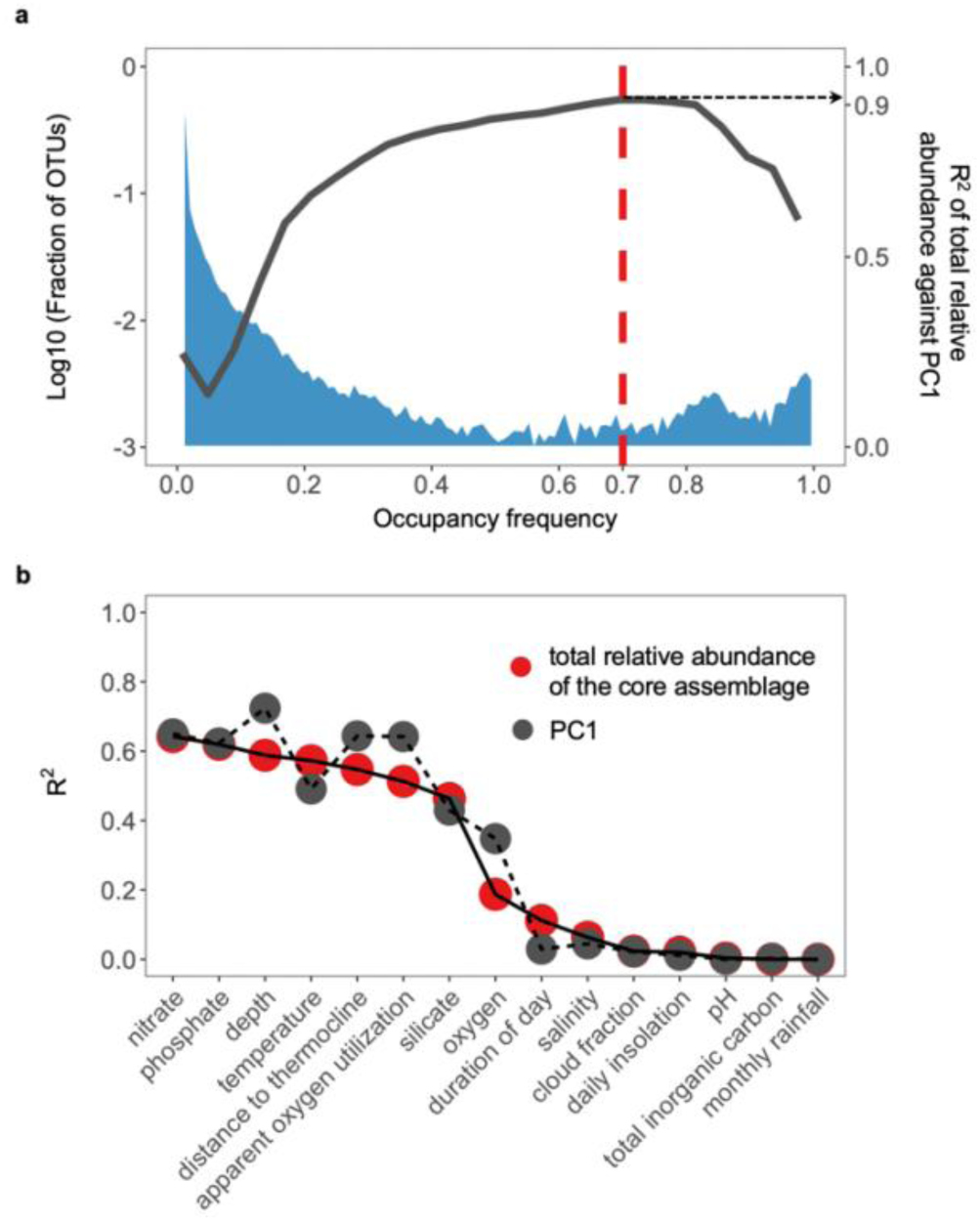
The relative abundance of the core assemblage, PC1 and a set of abiotic factors are strongly associated in the ocean microbiome. **a**, the distribution of occupancy frequency (the percentage of samples in which the taxa is present, blue, left y-axis) is u-shaped. A critical breakpoint is determined at 70%, as indicated by the first derivative of frequency function (Fig. S1). The highest linear regression R^2^ for total relative abundance of taxa against PC1 (black, right y-axis) is also obtained at >= 70% occupancy frequency cutoff. **b**, the relative abundance of the core assemblage is strongly correlated with a set of abiotic factors, such as nitrate concentration, temperature and depth.

Previous work with the Tara Ocean dataset showed a strong correlation between temperature and PC1 of the taxa-sample matrix^11^. Interestingly, we found that the core assemblage, made of the sum of those taxa with occupancy frequency no lower than 70%, was almost perfectly correlated to the PC1 of the ocean microbiome: the sum of core taxa explains 92% of the variation in PC1 (Fig. 1a). The principal component is a statistical device – a linear transformation of the taxa abundance that maximizes the variance explained. Although ordination axes of taxa-sample matrix such as principal components have been widely used in ecology to describe the association between community composition and environmental conditions^16,17^, their exact ecological or biological interpretation is not obvious. Our analysis shows that the PC1 actually captures the *core* ocean microbiome as defined by the critical point of the u-shaped occupancy frequency distribution.

The total relative abundance contributed by the core assemblage was not only strongly associated with PC1, but also with a set of abiotic parameters. Among them, the relationship between core assemblage relative abundance and nitrate concentration was the strongest (linear regression R^2^ = 0.64), followed by depth, temperature, phosphate concentration and apparent oxygen utilization (Fig. 1b). By comparison, random assemblages of taxa sampled from the whole microbiome with the same number of taxa as the core assemblage were only weakly associated with these environmental parameters (e.g. average R^2^ = 0.11 against nitrate for 1000 random samplings). Apart from this Tara Ocean dataset based on 16S miTags extracted from metagenomes, we further validated this result in an independent 16S amplicon-based ocean microbiome, the dataset of the ANT28-5 Latitudinal Transect of the Atlantic Ocean^12–14^. Here too we found a strong association between the relative abundance of the core assemblage, PC1 and temperature (linear regression R^2^ > 0.60, Fig. S2), with the core assemblage also defined at an occupancy frequency cutoff of 70%. One possible explanation for the observed variation in core assemblage frequency is that it is simply driven by changes in taxa richness, as opposed to a direct dependency between the abiotic factors and core taxa. Temperature is known to increase the richness of taxa in many environments^18^, potentially reducing the relative abundance of core taxa. However, we found that the association between taxa richness and temperature (linear regression R^2^ = 0.14) was much weaker than that between core assemblage and temperature (linear regression R^2^ = 0.57).

We hypothesize instead that the variation in the relative abundance of the core assemblage captures the biochemical structure of the ocean along a gradient of nutrient availability. Most of the parameters measured during the Tara expedition are not independent, but strongly correlated to one another. The concentrations of nitrate, phosphate and silicate are positively intercorrelated and they would all decrease as depth, temperature and apparent oxygen utilization increases (Fig. S3). All these parameters are thus different manifestations of the biogeochemical constraints associated with nutrient availability in the ocean. Therefore, we hypothesize that the core assemblage is composed of taxa that thrive in surface or pelagic waters, i.e. phototrophs and heterotrophs specialized for the uptake of dissolved organic carbon (DOC), whereas the flexible assemblage should be composed of organisms that thrive in deeper waters, such as those attached to sinking particles.

Consistent with this hypothesis, the core assemblage was enriched in SAR11 (standard effect size *SES*_*N*_ = 118.8), *Prochlorococcus* (*SES*_*N*_ = 34.4), *Synechococcus* (*SES*_*N*_ = 33.4), as well as The SAR86 clade within the order of *Oceanospirillales* (*SES*_*N*_ = 49.9) and Candidatus *Actinomarina* within the order of *Acidimicrobiales* (*SES*_*N*_ = 44.3) (Fig. 2a & Fig. 2b). By contrast, clades such as the *Alteromonadales*, frequently associated with particles in deeper waters^19^, were highly enriched in the flexible assemblage (*SES*_*N*_ = 11.9 in the flexible assemblage). Except for *Prochlorococcus* and *Synechococcus*, which are phototrophs, all the taxa highly enriched in the core assemblage are ‘oligotrophs’: highly specialized heterotrophic organisms, with a high affinity for DOC, slow growth rates and streamlined genomes^20–22^. In contrast, those taxa underrepresented in the core assemblage, such as *Alteromonadales, Flavobacteriales* and *Rhodobacterales* are metabolically versatile, fast growing organisms, typically associated to particles or phycospheres^23^. Therefore, we propose that the partitioning of the subtropical ocean microbiome into core and non-core assemblages captures a well-known fundamental divide in the ecological strategies of marine bacteria.

**Figure 2.**
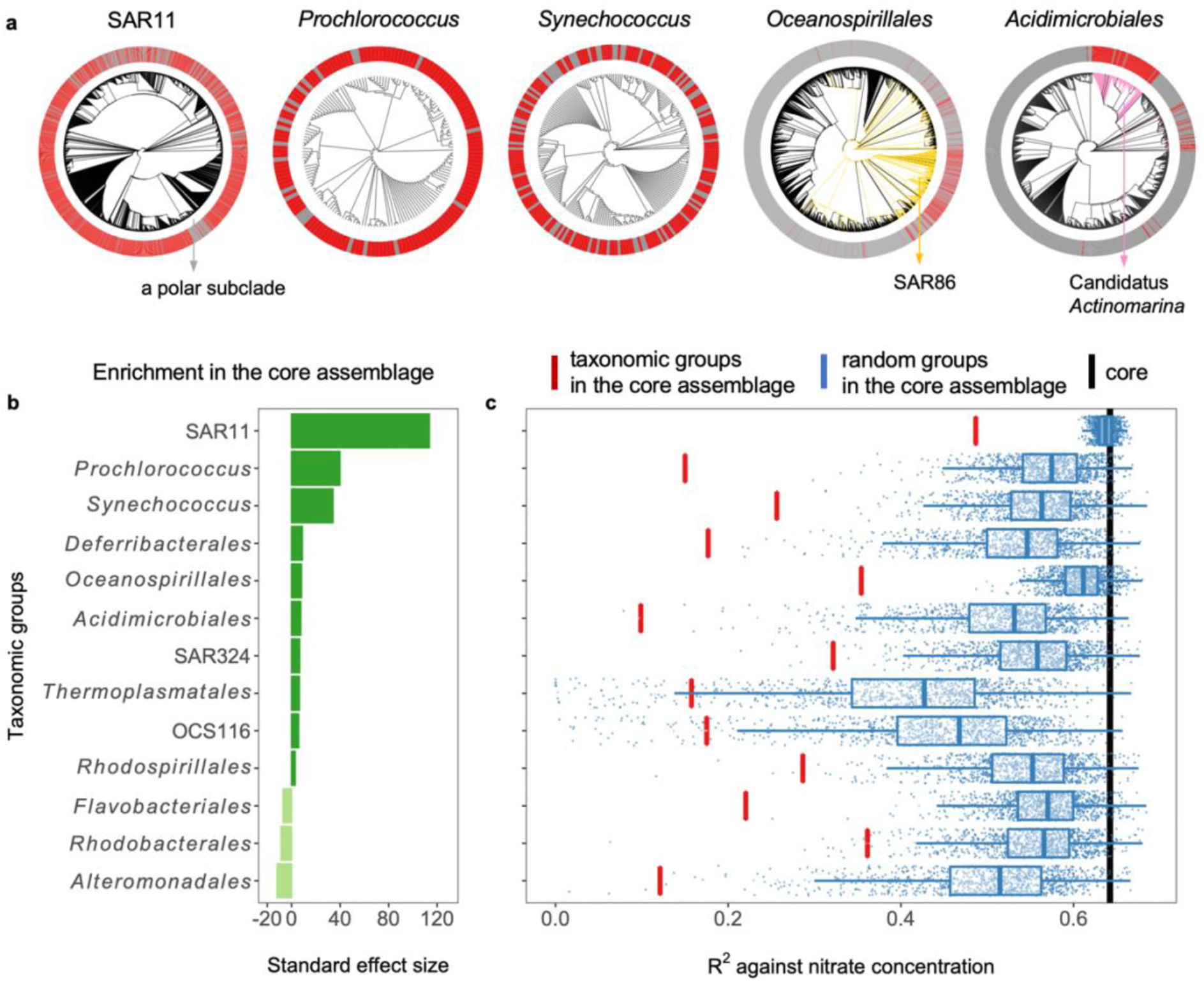
The taxonomic profile of the core assemblage. **a**. Phylogenetic signal of some representative clades highly enriched in the core assemblage. For each clade, core taxa are colored with red while flexible taxa are colored in grey. **b**. Standard effect size of enrichment for all the abundant taxonomic groups (average relative abundance > 1%) in the core assemblage, positive values (deep green) indicate enrichment in the core assemblage while negative values (light green) indicate enrichment in the flexible assemblage. **c**. The core assemblage as a whole (black) outperformed every individual taxonomic group (red) in terms of R^2^ against nitrate concentration. With the same number of taxa assembled across the core assemblage, random groups (blue) generally achieve higher R^2^ than taxonomic groups.

### The statistics of functional groups

The high percentage of variation in abiotic factors explained by the core as a whole could be caused by the grouping of a few taxa with strong association to those environmental parameters. However, we found this not to be the case. Instead, taxa in the core complemented each other to increase the R^2^ of the group against the environmental variables. We calculated the R^2^ of the linear regression for all the abundant taxonomic groups (average relative abundance > 1%) against nitrate as a focal abiotic factor. All constitutive taxonomic groups had lower correlations with the environmental parameters than the whole core assemblage (Fig. 2c). As a null model, we also calculated the R^2^ for random groups that are randomly assembled across the core assemblage. Interestingly, we found that with the same number of taxa, the R^2^ for taxonomic groups within the core assemblages were generally lower than that of random groups assembled across the core assemblage (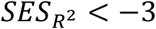, Fig. 2c). Thus, individual taxonomic groups not only did not drive the environment-core assemblage relationship, but underperformed relative to random groups of taxa. This means that the relationship between environment and taxonomic composition emerges from treating unrelated but complementary sets of organisms as one chimeric unit.

To understand this phenomenon, we developed a statistical framework, described in Box 1. In essence, the theory shows that there are two properties required for a taxa assemblage to have a higher correlation with an external parameter than any of its parts. We call those two properties *consistency* and *complementarity*. Consistency requires that the correlations between individual taxa and the abiotic factor should be strong and consistent in sign. More precisely, for each taxon, the projection of taxon vector on the environment vector should point in the same direction (Fig. 4, Box 1). Complementarity requires that, despite their consistency with respect to the abiotic factor, residual vector of taxa orthogonal to environment vector should be as negatively correlated with each other as possible (Fig. 4, Box 1). In this way, the errors of the residuals of the assemblage-environment fit contributed by each taxon will tend to cancel out, thus increasing the quality of the regression.

**Figure 3.**
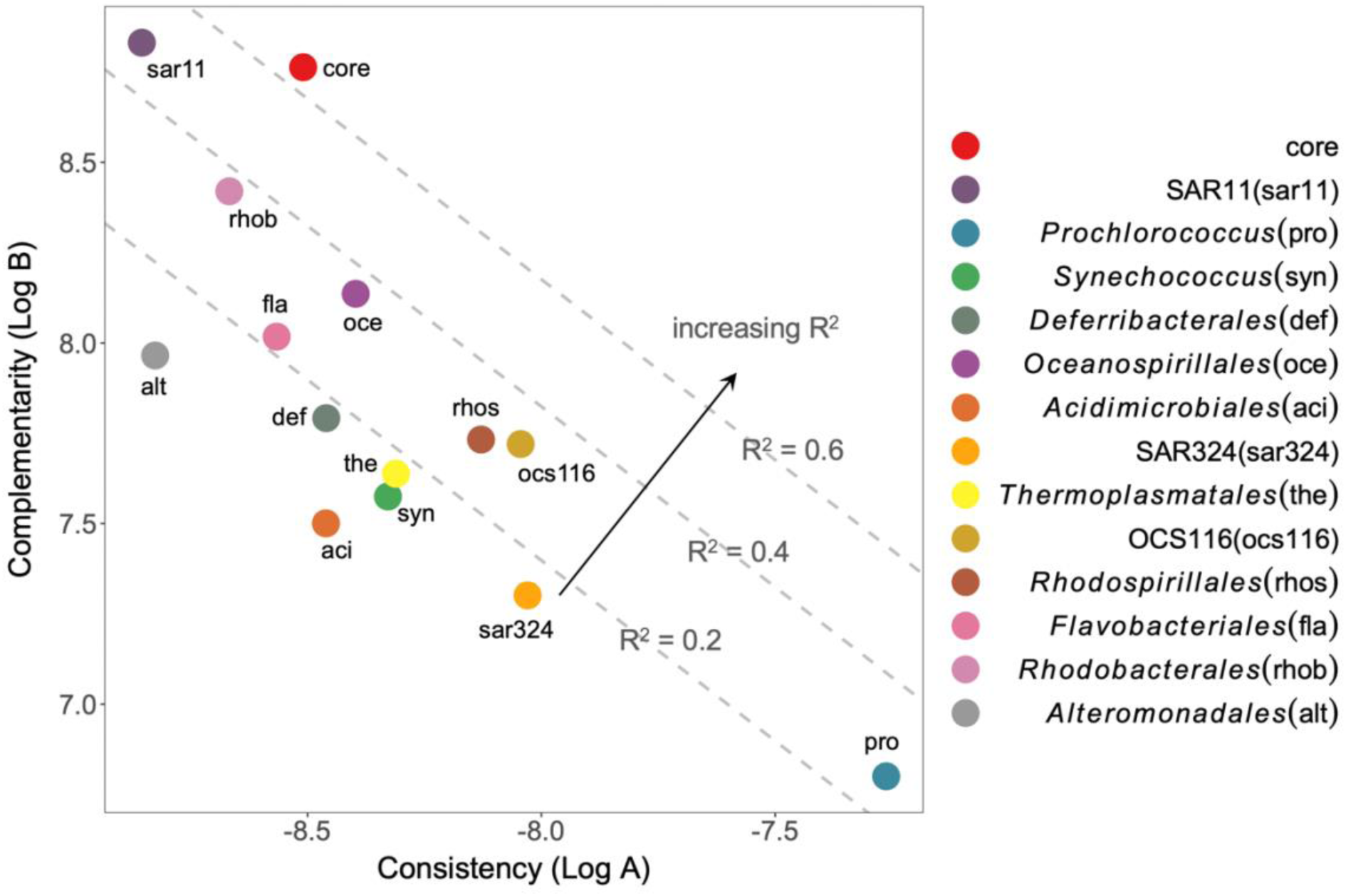
The interplay among consistency, complementarity and R^2^. In the log-scale Cartesian coordinates, moving upwards increases complementarity while moving rightwards increases consistency (Equation 4). Gray dashed lines with slope = −1 demonstrate “R^2^-cline” with different intercepts showing different R^2^ values. The direction of increasing R^2^ is thus perpendicular to R^2^-clines and points to the top right. Compared to most of the taxonomic groups, the core assemblage achieves far higher R^2^ since complementarity is improved at relatively small cost of consistency.

The statistical framework proposed in Box 1 helps us understand why taxonomic groups generally attained lower correlations with the abiotic factor than the core assemblage, or even random groups. The answer is that the taxa within a taxonomic group are on average more positively correlated to each other than randomly picked taxa, and thus exhibit lower complementarity. This means that within taxonomic groups the residuals of the taxa-environment regression are on average correlated, and thus get amplified when combining taxonomically related taxa. This, in turn, decreases the R^2^ of the regression. To visualize how each taxonomic group or assemblage behaved in terms of consistency and complementarity, we log-transformed Equation 1 (Box 1). This transformation allows us to obtain a linear relationship between log(A) and log(B), in which the intercept becomes a univariate monotonically increasing function of R^2^ (Equation 4). We found that thirteen out of fourteen abundant taxonomic groups exhibited lower complementarity compared to the core assemblage (Fig. 3). As these different taxonomic groups are added to the core assemblage, complementarity improves at the relatively small expense of consistency. The only taxonomic group with complementarity higher than the core assemblage was the SAR11 clade, implying that different SAR11 OTUs replace each other across different geographic locations^24^. On the other end, *Prochlorococcus* was the taxonomic group with the lowest complementarity, implying that all members of this clade tend to change in abundance across the ocean in a coordinated fashion.

#### Box 1 Consistency and complementarity together shape a functional group.

**Figure 4.**
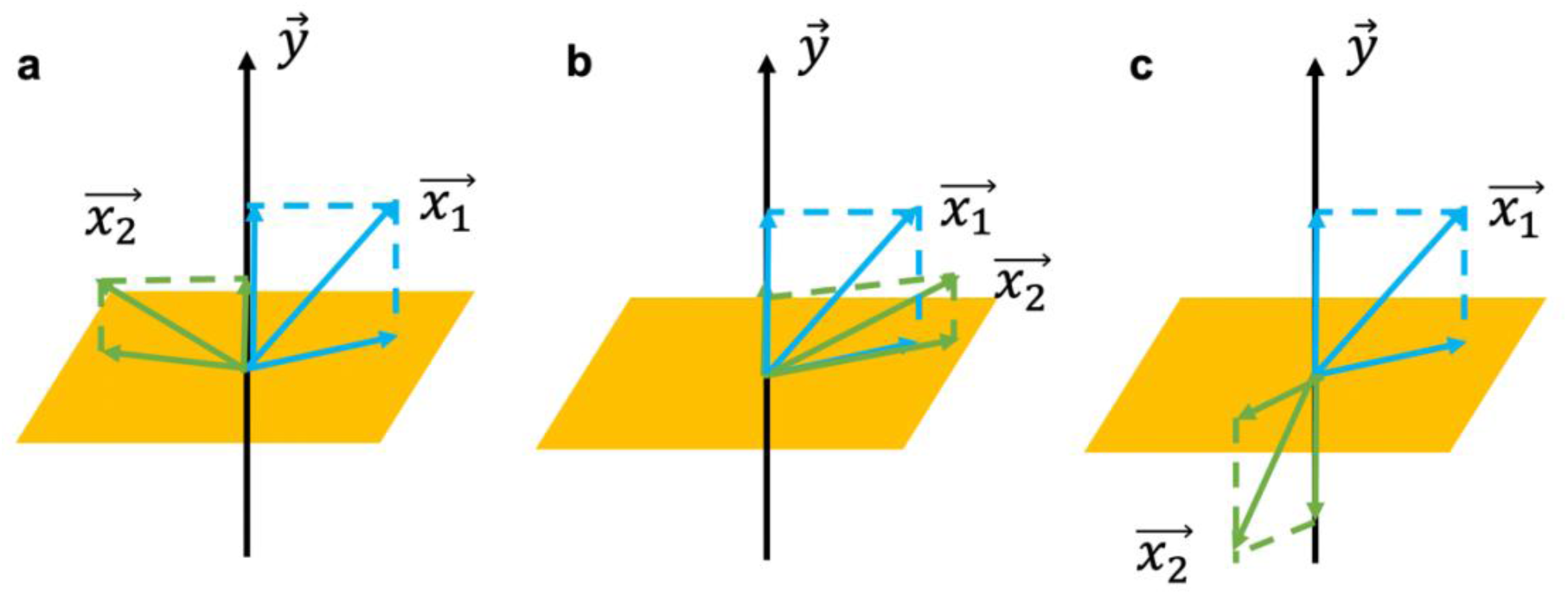
Schematic view of mathematical framework of consistency and complementarity for Box 1. In this figure we consider the simplest two-dimensional case, where a taxa assemblage 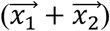 is to correlate with an abiotic factor 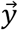. Every taxa vector could be decomposed into two components, one parallel to 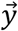 and the other in the subspace orthogonal to 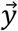. The R^2^ will be maximized if 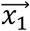 and 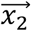 are consistent in the parallel direction and complementary in the orthogonal subspace (**a**). Compared to **a, b** breaks the condition of complementarity and **c** breaks the condition of consistency.

One of the desirable properties of a meaningful grouping of taxa is for it to predict abiotic parameters better than any of its constituent taxa. But what are the conditions that make this happen? To answer this question, we take a look at the math behind a taxa assemblage and correlations. Suppose ***x***_*i*,_ *i* ∈ {1,2, …, *n*} is the vector of relative abundances at different sampling locations for the i-th taxon, and ***y*** is the corresponding vector for an abiotic factor, e.g. nitrate concentration. Then the vector of relative abundances for a taxa assemblage, e.g. a functional group, is given as 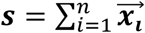. This sum of taxa vectors is actually implemented in two orthogonal directions (Fig. 4). One is in the direction parallel to the vector corresponding to the abiotic factor (i.e. the projection on ***y***), while the other is in the subspace orthogonal to the abiotic factor (i.e. the residuals). In order to optimize the correlation between the vectors ***s*** and ***y***, the sum of projection vectors should be maximized, while the sum of residual vectors should be minimized. In other words, taxa within a functional group are required to be consistent in the parallel direction (i.e. consistency) but complementary in the orthogonal subspace (i.e. complementarity). Consistency indicates that taxa have similar responses to the abiotic factor shaping that functional group. Complementarity implies competition or ecological differentiation in other independent niche axes^1^.

Consistency and complementarity can be captured by a simple mathematical expression (see Methods for full details). As in the previous sections, we took the R^2^ of the linear regression between a functional group and an abiotic factor as a measure of the strength of their association. We found that the R^2^ for a functional group ***s*** and an abiotic factor 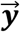 can be decomposed as,

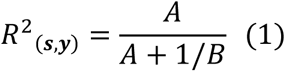

where

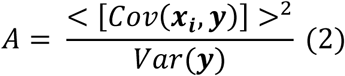

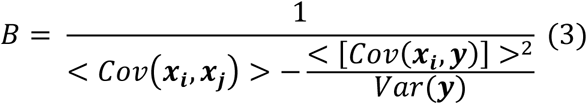

*A* is the square of average taxa-environment variance for a functional group, scaled by the variance of the abiotic factor. Larger values of *A* indicate consistent response to abiotic factors among taxa. *B* is the inverse of the difference between average taxa-taxa covariance and average taxa-environment covariance, which can be regarded as a “partial” average taxa-taxa covariance after controlling the abiotic factor. Higher values of B indicate higher complementarity between taxa.

A logarithmic transformation allows us to obtain a linear relationship between log(*A*) and log(*B*), in which the intercept is univariate function of R^2^.

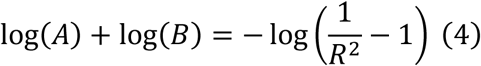

Note that on the right-hand side it is a monotonically increasing function with *R*^2^. In this sense, a high *R*^2^ is jointly contributed by a larger value of log(*A*) and a larger value of log(*B*), which are quantitative indicators for higher consistency and higher complementarity.

### Most supervised functional groups are skewed in the core-flexible bipartition

The statistical framework developed above and the strong partitioning into core and non-core assemblages revealed in the microbiome give us the tools to understand how other taxa aggregations behave in relation to abiotic factors. In particular, we focused on a previous study by Louca et al. that showed that groups of taxa with common metabolic annotation displayed much stronger associations with abiotic factors than individual taxa or taxonomic groups^5^. These groups were proposed to represent functional groups, created in a supervised manner by linking taxa names to central metabolic functions like denitrification, methanotrophy, sulfate reduction, etc. We found that compared to the whole community, 27 out of 28 of these supervised functional groups were more skewed in their abundance composition in terms of core vs. flexible bipartition (Fig. 5a). For instance, 95.0% of the reads in the supervised group ‘photoautotrophy’ were contributed by members of the core assemblage, e.g. *Prochlorococcus* and *Synechococcus*. Conversely, the group ‘cellulolysis’ was exclusively composed of flexible taxa.

**Figure 5.**
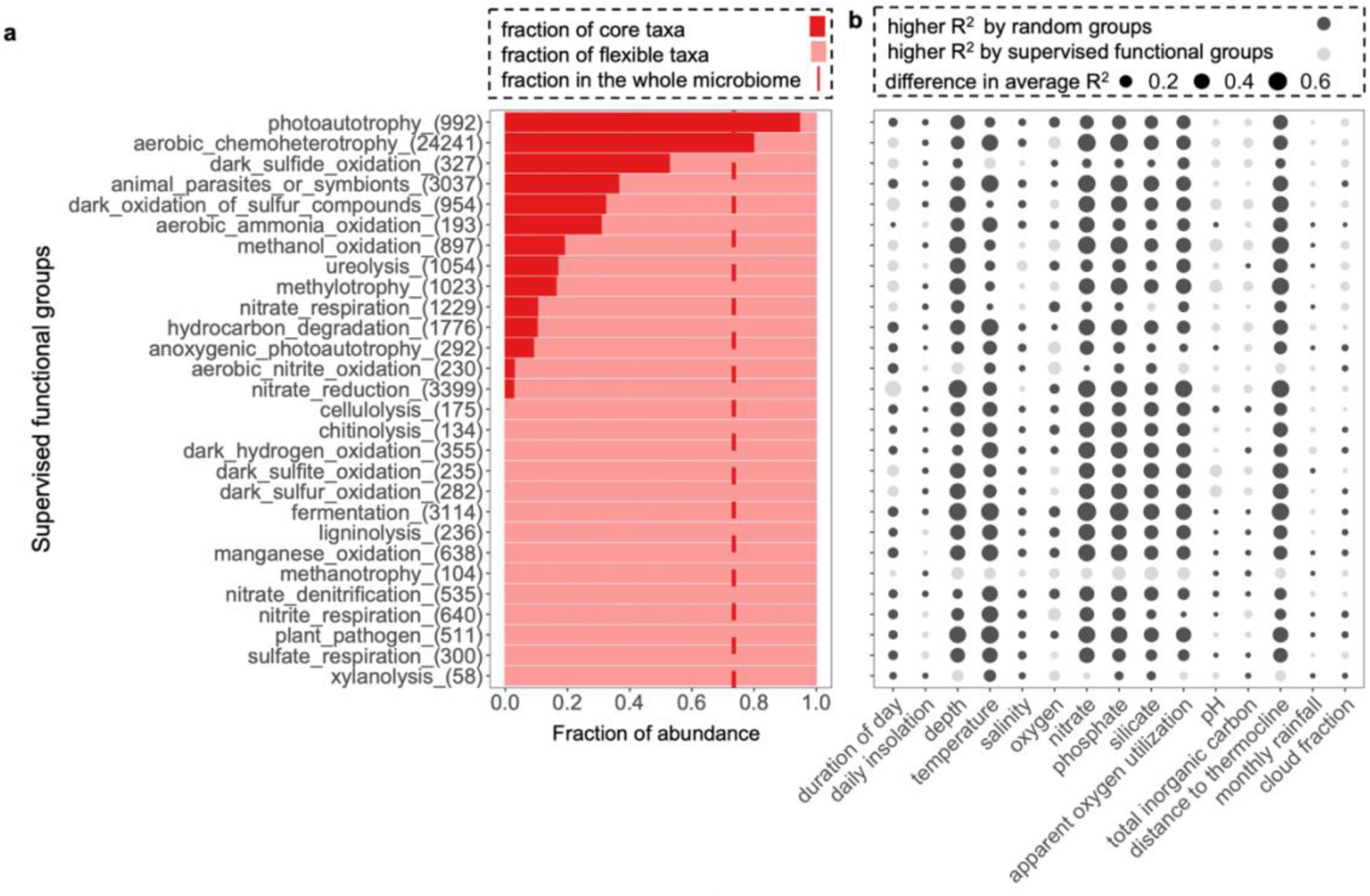
The strong environmental association of supervised functional groups may result from the fact that most of them are nested in either the core or flexible assemblage. **a**. Compared to the whole microbiome (red dashed line, 73.5% of core taxa and 26.5% of flexible taxa), most functional groups are more skewed in their core (deep red) vs. flexible (pale red) composition. The number of OTUs in each functional group is in the bracket after their names. **b**. For each supervised functional group, we assemble random groups by randomly sampling taxa from the whole microbiome based on the same bipartite composition of core vs. flexible taxa. For environmental parameters such as nitrate, depth, temperature and apparent oxygen utilization, random groups generally outperform supervised groups in terms of R^2^, while for pH and total inorganic carbon supervised groups generally attain higher R^2^.

The predominance of either core or flexible taxa in these annotation-based functional groups raises a possibility that their strong environmental association might just be a consequence of being nested within the core or flexible assemblage. To test this hypothesis, we regressed the environmental parameters against random groups with the same number of taxa and same core vs. flexible composition as each supervised functional group. We found that, in terms of their correlation with nitrate, depth, temperature, phosphate, silicate and apparent oxygen utilization, random groups generally outperformed supervised functional groups, which were less phylogenetically diverse than their random counterparts (Fig. 5b & Figure S4). This analysis suggests that being nested in the either core or flexible assemblage, where taxa are on average consistent and complementary, is sufficient to explain the strong environmental association observed for supervised functional groups. However, for abiotic factors such as pH and total inorganic carbon, supervised functional groups displayed stronger correlations than random groups, suggesting that CO_2_ concentration and solubility in seawater may influence the abundance of more specific subsets of taxa.

## Discussion

Our analysis shows that a bipartition into core and flexible assemblages of taxa captured a fundamental aspect of the structure of the ocean microbiome, i.e., the biogeochemical constraints imposed by gradients of nutrient availability. ‘Core’ taxa, composed of cyanobacterial phototrophs and oligotrophic heterotrophs with streamlined genomes, such as SAR11, *Actinomarina* and SAR86, dominate in surface waters or pelagic waters and their distribution is almost perfectly captured by PC1 of the Tara Oceans microbiome. In contrast, a more diverse and metabolically versatile set of taxa, such as *Alteromonadales, Flavobacteriales* and *Rhodobacterales* (e.g. *Rueggeria*), increase in relative abundance with nutrient availability and depth. This simple pattern that we derived from the combination of taxa distributions and abiotic factors captured by the Tara expedition is the main structure-function feature of the global ocean microbiome. In concordance with this idea, we found that the statistical association between supervised functional groups - based on annotations - and abiotic factors is largely driven by the fact that these functional groups are nested within core and flexible assemblages. A null model in which groups are created by randomly picking taxa based on the same bipartite composition reproduces most of the statistical associations observed with supervised functional groups. This does not mean necessarily that those supervised functional groups are not meaningful, but simply that they cannot be independently validated based on the current data. Doing so would require a larger number of independent environmental parameters.

A functionally meaningful grouping of taxa should, in theory, be better-correlated with the abiotic parameters that constrain its function than individual taxa. In this study we found the conditions required to achieve such higher correlations when grouping taxa into functional groups. We call these conditions consistency and complementarity. Consistency requires members of a functional group to be consistent in their association with the abiotic factor, even if the strength of this association is near zero. This is similar to the concept “functional response group” proposed in a previous study^25^, which means that taxa respond similarly to certain abiotic factor. However, that concept does not consider the relationship between taxa within an assemblage. In our framework, the relationship between taxa is quantified by complementarity. A high degree of complementarity implies that the grouped taxa exclude each other at different locations, possibly due to competition for a common resource and being better adapted to the local environmental conditions.

The development of algorithms that detect functional groups from patterns of taxa abundance and abiotic factors, independently of external annotations, remains an open challenge. Such algorithms could rely on new experimental designs, tailored to the question. For instance, studying the response of the different taxa to pulse perturbations, (e.g. externally supplied resources, temperature changes, salinity, etc.) could help us identify groups of taxonomically distinct organisms that respond to environmental changes in a similar fashion. This type of data, along with patterns of taxa variance and covariance across global datasets could open new avenues to identify the functional units of microbiomes.

## Methods

### Mathematical framework

Let ***x***_*i*,_ *i* ∈ {1,2, …, *n*} be vectors for n taxa within an assemblage and ***y*** be the vector for an abiotic factor, e.g. nitrate concentration in this paper. The length of ***x***_*i*,_ *i* ∈ {1,2, …, *n*} and ***y*** are given by the number of samples in the dataset, *p*. For instance, the k-th element of ***x***_*i*_, ***x***_*ik*_, denotes the relative abundance of taxon *i* in sample *k*. Similarly, the k-th element of ***y***, ***y***_*k*_, denotes the nitrate concentration in sample *k*. For simplicity, ***x***_*i*_ and ***y*** can be centered to zero mean without altering their correlations.

Let ***s*** be a column vector of length *p*, representing the total relative abundance of the taxa assemblage across the *p* samples. We can measure the strength of the relationship between ***s*** and ***y*** by calculating the coefficient of determination, or R^2^, which in statistics is equal to the square of the Pearson’s correlation coefficient. Pearson’s correlation coefficient, by definition, is the covariance weighted by the products of standard deviation. For vectors centered at zero, this is the same as the inner product of vectors weighted by the product of vector lengths, which is actually the cosine of the angel between two vectors. *r* = *cosθ*.

We can decompose our vector of taxa assemblage ***s*** into two components (Fig. 4), with ***a*** being the vector component that is parallel to ***y*** (in other words, the projection of ***s*** on ***y***), and ***b*** being the component in the subspace that is orthogonal to ***y*** (in other words, the residual of projection). In this way we can rewrite *R*^2^ using only ***a*** and ***b*** as

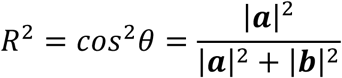

With the formula of projection, we have

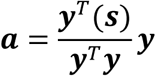

Thus

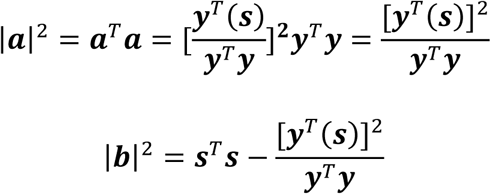

It would be operationally much easier to use this framework if the algebraic equations could be translated into a simpler expression. To simplify, we can rewrite |***a***|^2^ and |***a***|^2^ **+** |***b***|^2^ as

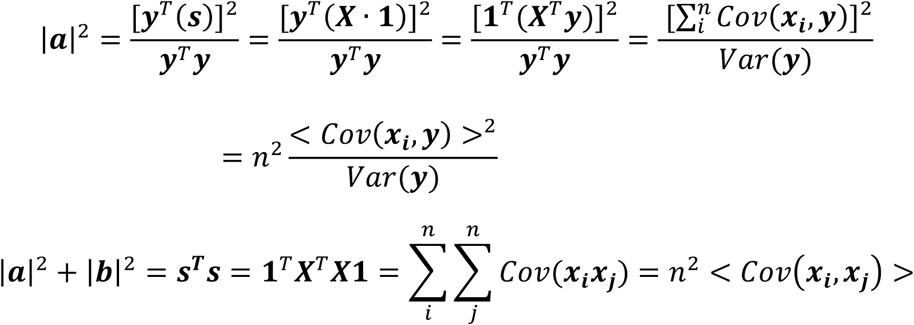

Where **1** is a column vector of length p and all elements equal to 1. Therefore, the R^2^ for an assemblage of taxa linearly regressed against an abiotic factor is

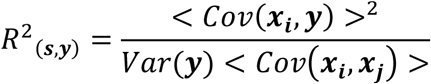

A convenient transformation that allows us to disentangle the effects of complementarity and consistency is

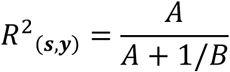

where

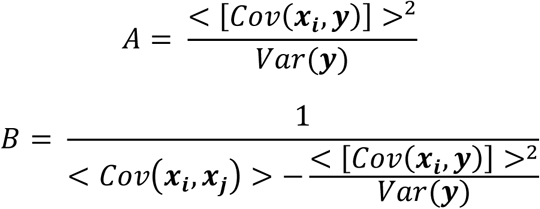

Larger values of *A* denote higher consistency, implying the taxa within an assemblage have correlations of the same sign with the abiotic factor. Larger values of B implying high complementarity, implying that taxa the residues of the taxa vs. abiotic factor regression tend to be anti-correlated with each other. Notice the denominator in the expression for B is always a positive number: By the properties of covariance matrices, even if the inter-taxa covariance is negative, the average over the whole covariance matrix is positive.

### Sequencing data

The Tara Ocean microbiome dataset used in this study is exactly the same as that used in previous research^5^. In brief, the quality-filtered DNA sequences of the 16S rRNA gene were extracted from metagenomes^11^ and were clustered to generate OTUs by closed-reference mapping to SILVA v119 database at 99% similarity^5^. There are in total 43812 OTUs for 124 samples, accompanied by a metadata table of 15 environmental parameters including temperature, depth, nitrate concentration, etc. With the representative sequences for OTU, we constructed a phylogenetic tree based on SATe-enabled phylogenetic placement (SEPP)^26^. We also repeated the analyses using the newer SILVA v132 database^27^ but the results remained consistent. We therefore used SILVA v119 for its better compatibility with the FAPROTAX software for annotation-based functional group calling5. In addition to the Tara Ocean dataset, we also used an independent dataset, the ANT28-5 Latitudinal Transect of the Atlantic Ocean to validate the relationship between PC1, the core assemblage and abiotic factors such as temperature. This dataset was shared via Simons Collaborative Marine Atlas Project (CMAP). In brief, the publicly accessible 16S rRNA amplicon reads were merged and quality-filtered with VSEARCH and then amplicon sequence variants (ASVs) were generated with Deblur^12–14^. There were in total 4873 ASVs for 130 samples, with a metadata table for temperature, depth and salinity.

### Statistical analyses

In order to quantify the level of enrichment for different taxa in the core assemblage, we calculated the standard effect size as

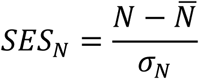

*N* is the number of OTUs for a certain taxonomic group in the core assemblage, 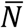 and *σ*_*N*_ are the mean and standard deviation of *N* for a null model in which 999 random assemblages with the same number of taxa as the core assemblage were subsampled from the whole microbiome.

Similarly, we also calculated the standard effect size of R^2^ as

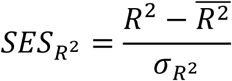

where *R*^2^ is the linear regression *R*^2^ for a certain taxonomic group against nitrate concentration, 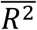 and 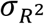 are the mean and standard deviation of *R*^2^ for a null model in which 999 random assemblages with the same number of taxa as that taxonomic group were subsampled from the core assemblage.

### Other statistical analyses

All the other statistical calculations, including correlations, covariances and linear regressions were performed in R^28^. Partial correlations were calculated with the package of *ppcor*^29^. All visualizations were performed using the package *ggplot2*^30^ and the phylogeny was visualized using the package *ggtree*^31^.

## Acknowledgements

The authors thank Stilianos Louca and Jesse McNichol for sharing the raw data, Morgan Ludwig and Lawrence Gardner for helping with the computing cluster as well as Cameron Thrash for helpful suggestions. We also thank all members of the Cordero lab, in particular Matti Gralka, Leonora Bittleston, Rachel Soble, Gabriel E. Leventhal and Shaul Pollak, for their support and useful discussions. OXC was supported by Simons Early Career Award 410104 and the Simons Collaboration: Principles of Microbial Ecosystems, award number 542395. XS and OXC were supported by NSF-BSF grant DEB 1655983.

## Author contributions

X.S. and O.X.C designed the study, analyzed the results and wrote the paper.

## Supplementary Figures

**Figure S1.**
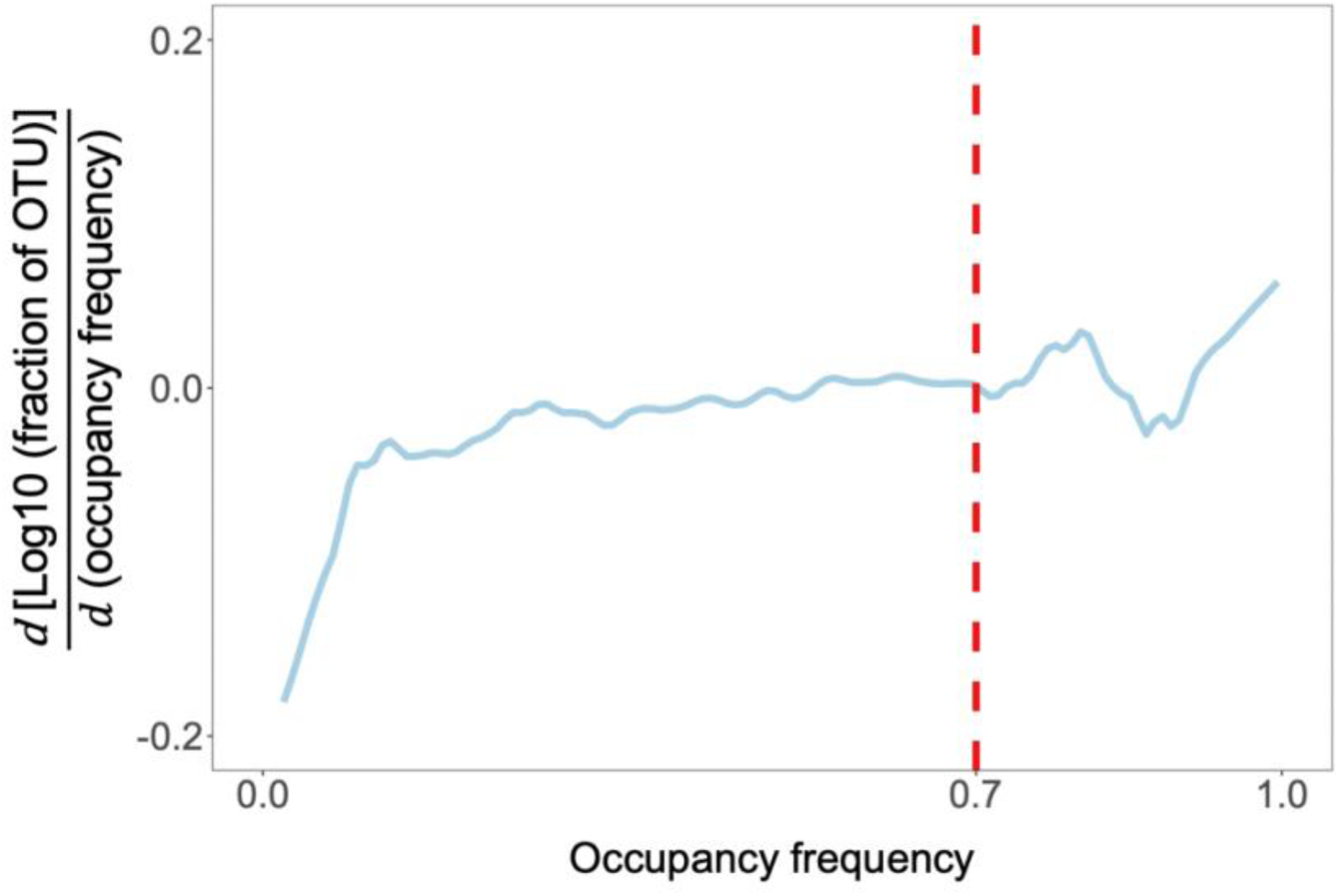
The critical point for the u-shaped occupancy frequency distribution is determined at 70% where the frequency function starts to increase rapidly, as indicated by a bump in the first derivative of frequency function.

**Figure S2.**
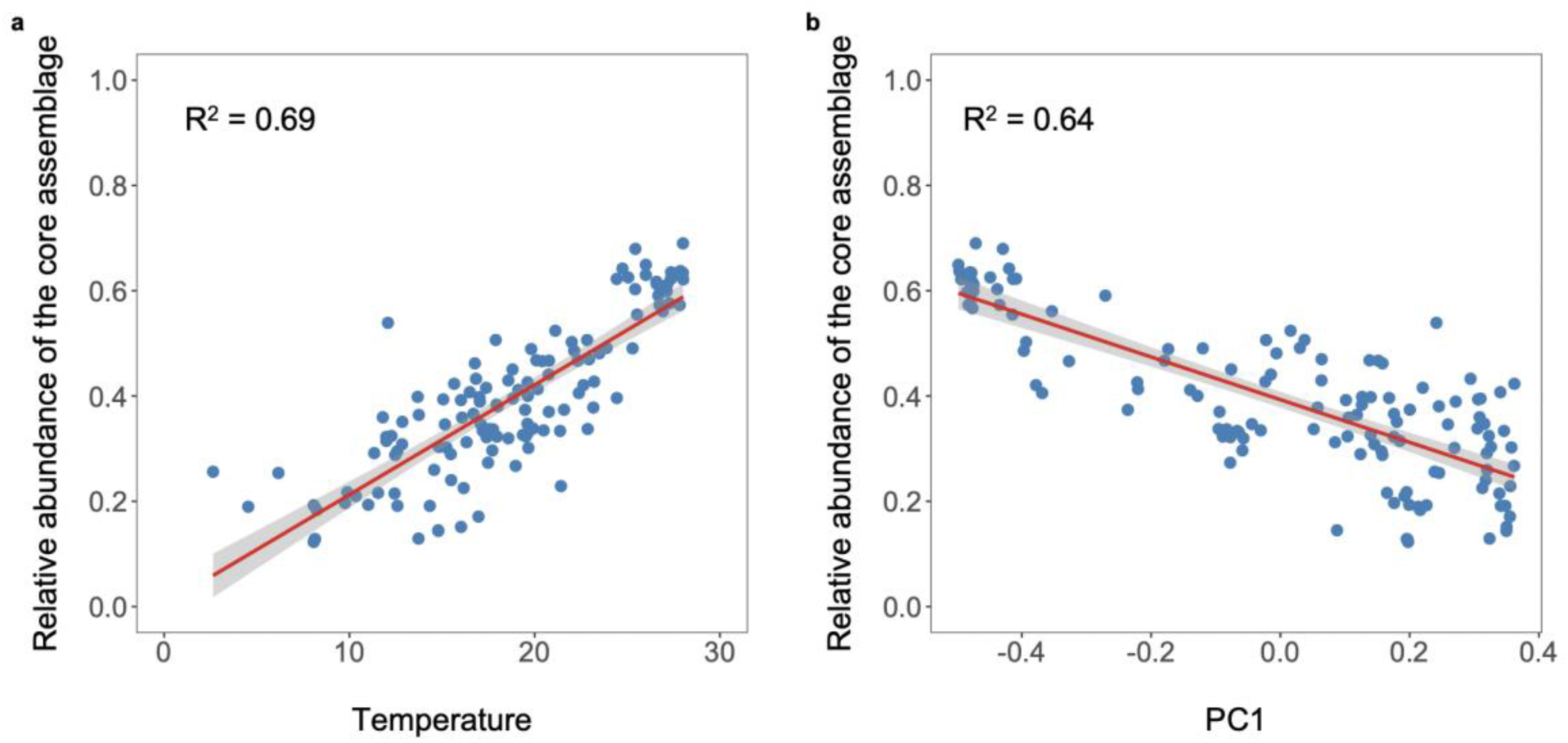
Relative abundance of the core assemblage, PC1 and temperature are also strongly associated in another independent dataset, the ANT28-5 Latitudinal Transect of the Atlantic Ocean. The core assemblage in this dataset is also assembled by taxa with >70% occupancy frequency.

**Figure S3.**
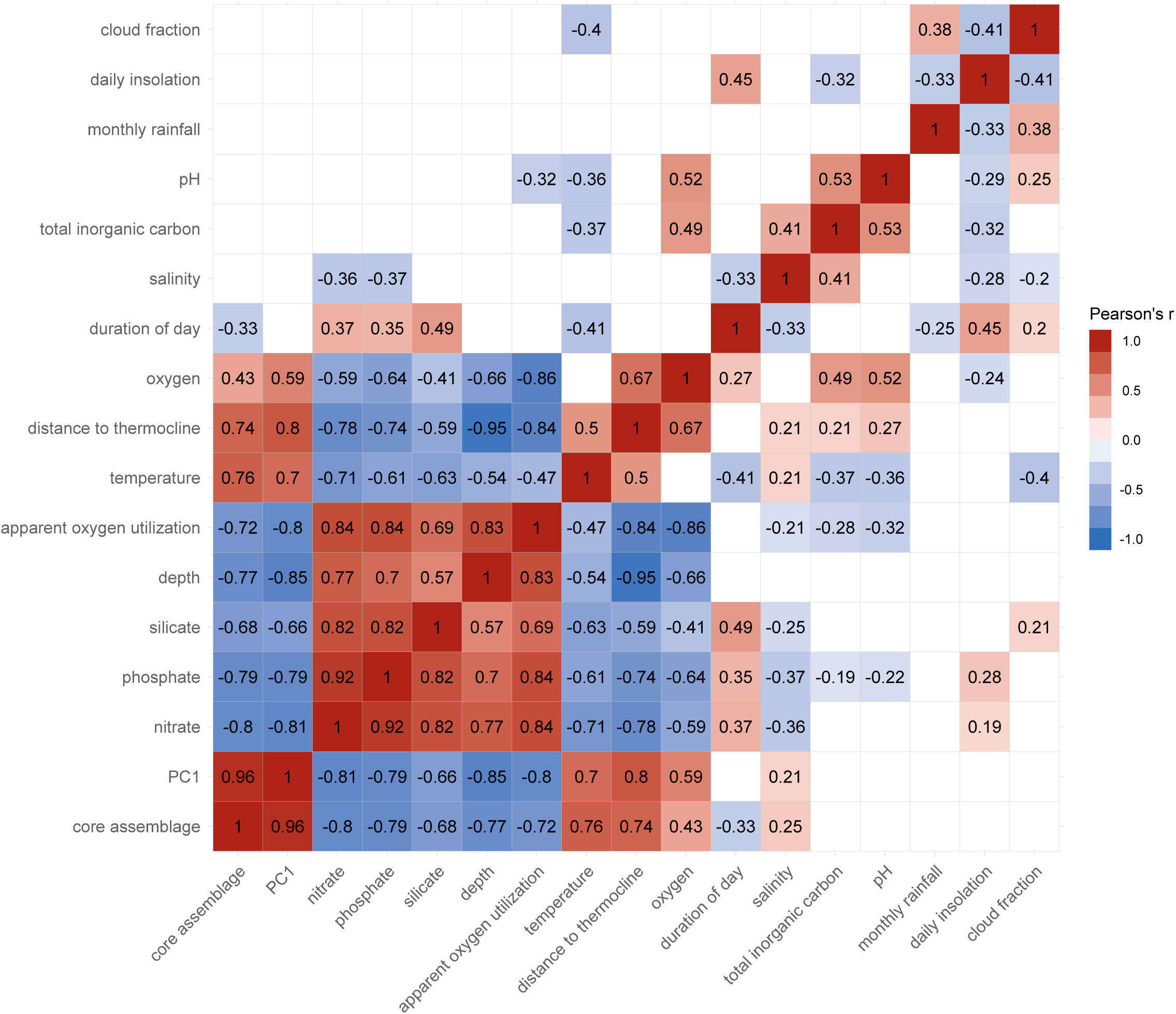
Intercorrelation between the core assemblage, PC1 and all the abiotic factors. Only significant correlations (*P* < 0.05) are shown in color tilts.

**Figure S4.**
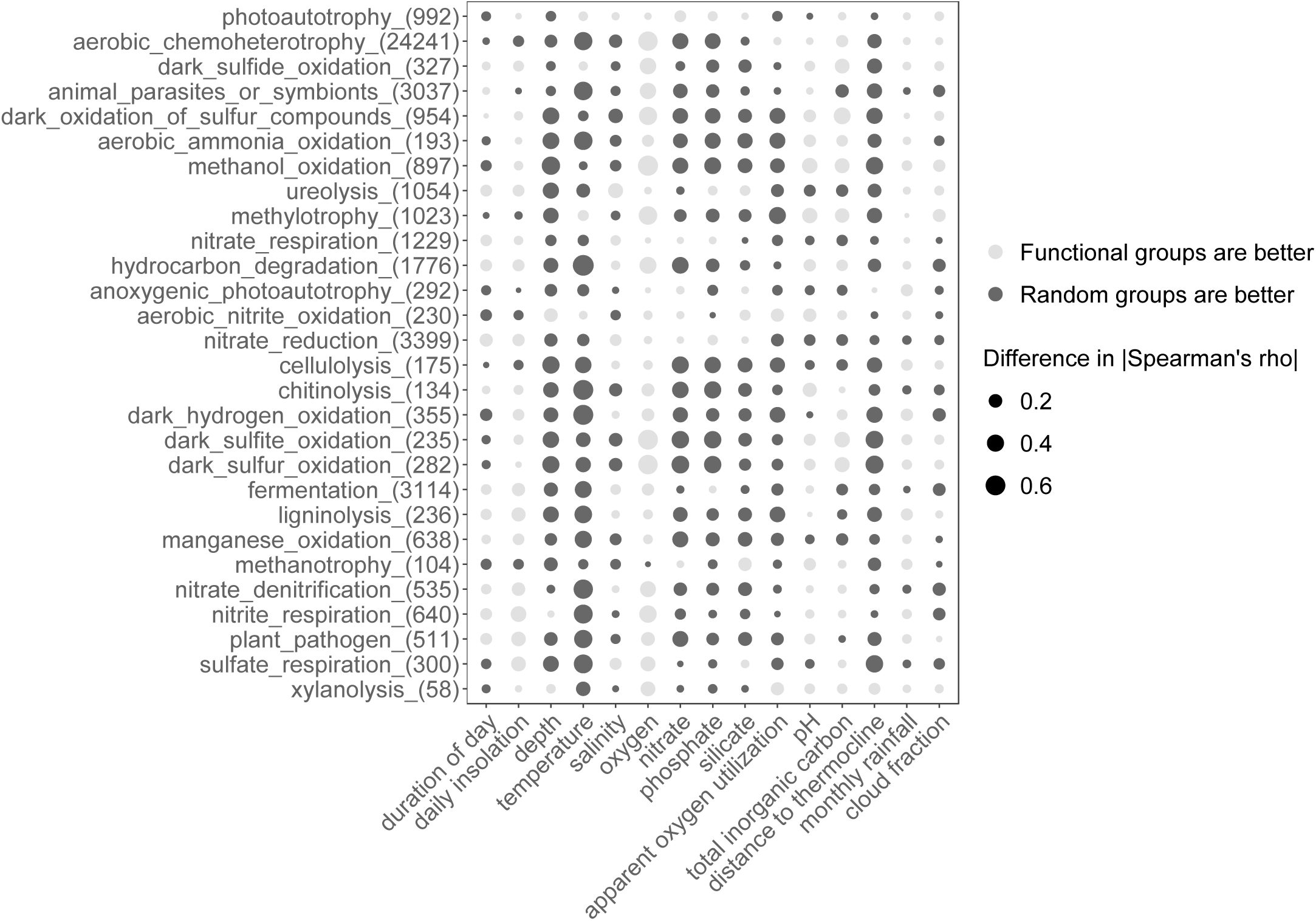
For abiotic factors such as nitrate, depth, temperature and apparent oxygen utilization, random groups generally outperform supervised groups in terms of spearman’s rho, while for pH and total inorganic carbon supervised groups generally attain higher rho.

